# Evidence for metacognitive bias in perception of voluntary action

**DOI:** 10.1101/423244

**Authors:** Lucie Charles, Camille Chardin, Patrick Haggard

**Affiliations:** Institute of Cognitive Neuroscience, University College London, London WC1N 3AR United Kingdom; École Normale Supérieure, Département des Études Cognitives, Paris

**Keywords:** Action, Metacognition, Confidence, Volition

## Abstract

Studies of metacognition often measure confidence in perceptual decisions. Much less is known about metacognition of action, and specifically about how people estimate the success of their own actions. In the present study, we compare metacognitive abilities between voluntary actions, passive movements matched to those actions, and purely visual signals. Participants reported their confidence in judging whether a brief visual probe appeared ahead or behind of their finger during simple flexion/extension movement. The finger could be moved voluntarily, or could be moved passively by a robot replaying their own previous movements. In a third condition, participants did not move, but a visual cursor replayed their previous voluntary movements. Metacognitive *sensitivity* was comparable when judging active movements, during passive finger displacement and visual cursor reply. However, a progressive metacognitive *bias* towards overconfidence was found for passive and for voluntary movements. Taken together, our results may partly explain some of the peculiarities that arise when one judges one’s own actions.

## 1 Introduction

What do humans know about their motor actions? And can they judge accurately their own movements?

A key feature of our cognitive system is the ability to monitor the accuracy of its own processing, a cognitive function generally described as metacognition (Fleming & Frith, 2014). This ability translates into a degree of confidence associated with each of our actions and decisions. It remains debated whether metacognitive judgments in different tasks rely on distinct specialized cognitive modules specific to each task or rather depend on a common single metacognitive function. Since the ability to judge our performance strongly depends on how good we are at performing a task in the first place (Galvin, Podd, Drga, & Whitmore, 2003; Maniscalco & Lau, 2012, 2016), comparing ‘second-level’ metacognitive abilities across tasks requires careful control for ‘first-level’ task performance. A statistical model of the relationship between first and second-order decisions offers a formal way to do this. This method has suggested that metacognitive function can be specifically impaired independently of decision accuracy (Rounis, Maniscalco, Rothwell, Passingham, & Lau, 2010). Prefrontal regions are associated with this ability (Fleming, Weil, Nagy, Dolan, & Rees, 2010), suggesting a supra-modal general-purpose centre for metacognition. Neuroimaging data further shows that different types of motor errors evoked a similar neural signal for incorrect actions (Falkenstein, Hoormann, Christ, & Hohnsbein, 2000; Gehring & Fencsik, 2001). Such findings suggest that confidence could provide a common currency for the brain to compare the accuracy of different types of decisions (Ais, Zylberberg, Barttfeld, & Sigman, 2016; De Gardelle, Le Corre, & Mamassian, 2016).

Most studies on metacognition involve first-level judgements of visual or auditory stimuli (see Faivre, Filevich, Solovey, Kühn, & Blanke, 2017 for an exception). It remains unclear however whether metacognition for interoceptive and proprioceptive signals differs from metacognition for visual and auditory stimuli (Garfinkel, Seth, Barrett, Suzuki, & Critchley, 2015). On the one hand, one could argue that humans have better metacognitive representations of their own movements than of external events. This argument is based on privileged access to information about our own self (Hart, 1965; Metcalfe & Greene, 2007). Indeed, knowing with precision the degree of certainty about limb position and bodily state is crucial for our survival and one could hypothesize that therefore we have better metacognitive access to these types of information than for any other type of signal. On the other hand, experimental data seem to suggest that humans have surprisingly first-level awareness about their own actions and somatic states (Garfinkel et al., 2015) It has been shown for instance that humans have relatively low accuracy in proprioceptive judgment, since strong illusions regarding limb position or body ownership are readily created by altering visual feedback (Blanke, Slater, & Serino, 2015). Fourneret & Jeannerod confirmed that participants could remain dramatically unaware of well-organized movement adjustments (Fourneret & Jeannerod, 1998). Similarly, it has been shown that humans have only limited awareness of some type of actions such as eye movement programs (Endrass, Reuter, & Kathmann, 2007; Nieuwenhuis, Richard Ridderinkhof, Blom, Band, & Kok, 2001; van Zoest & Donk, 2010). These results fit with the view that coordinated motor behaviours are often controlled unconsciously by specialized spinal and cerebellar circuits operating outside of awareness. This has led some authors to propose that motor awareness is confined to initiation of actions and evaluation of outcomes, with only limited access to motor commands themselves (Blakemore & Frith, 2003; Blakemore, Wolpert, & Frith, 2002). In sum, we normally know that we prepare and initiate action, and we know from sensory feedback whether our actions produce the intended outcome or not, but we have little access to the details of voluntary movements themselves (Haggard, 2017).

To our knowledge, no study has formally investigated metacognition for one’s own actions. In the present study, we investigated whether metacognitive abilities for perception of voluntary movements, differed from those for perception of either kinematically-matched passive movements or moving visual stimuli. To do this, we used a robotic device capable of recording finger position during active movements, as well as move the finger passively. In a novel dynamic position sense task, participants judged the instantaneous position of their flexing and extending finger (or of a moving dot in the visual only condition) relative to the position of a probe which unpredictably appeared on a screen along with the visual cursor displaying current finger position (Figure 1). By contrasting confidence judgments on Active finger movements, Passive finger displacement and Visual replay of the movements we were able to test the hypothesis of privileged metacognitive access to voluntary actions.

**Figure 1:**
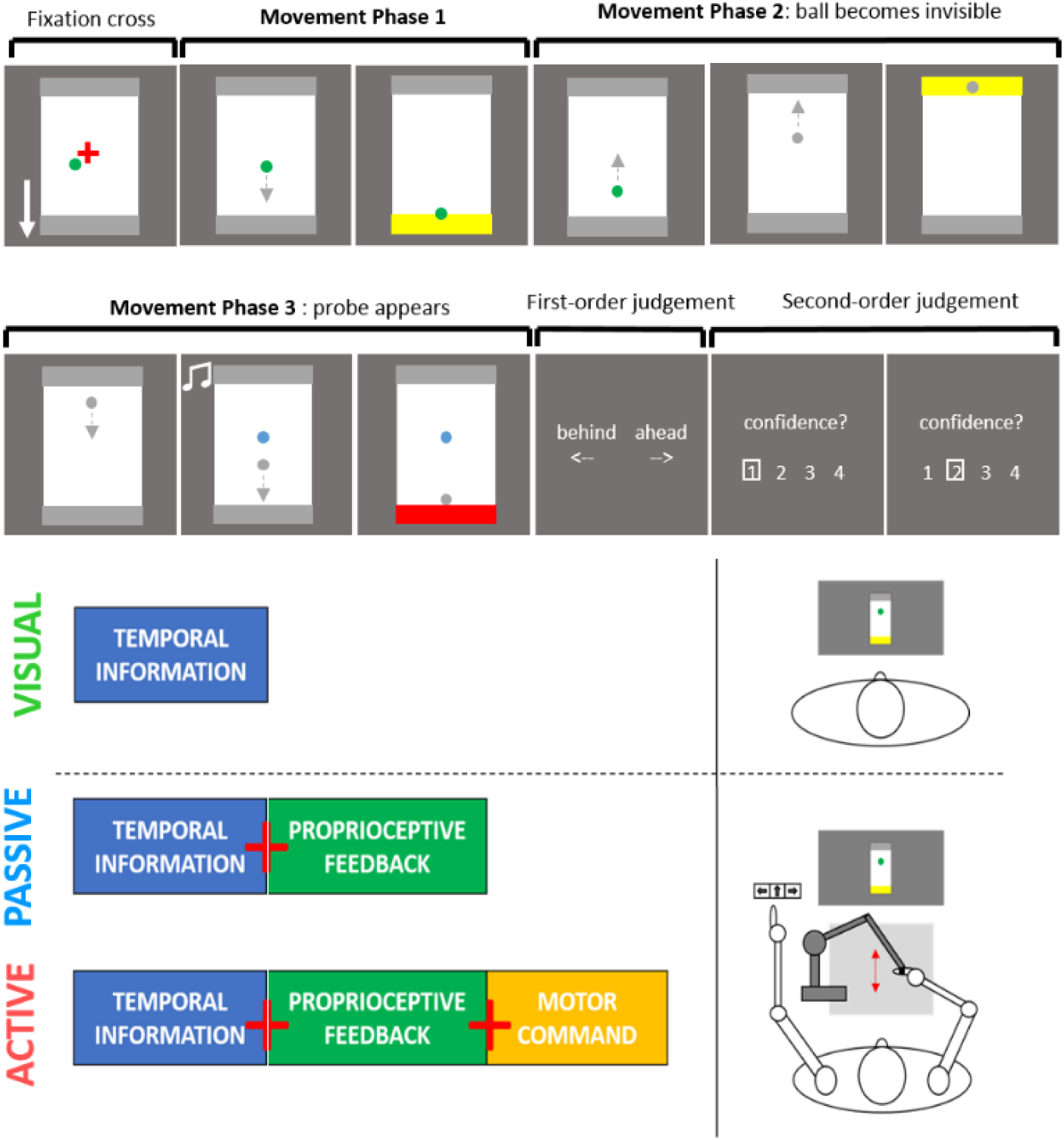
Protocol and design of the experiment. The participant’s finger was attached to a robotic arm capable of recording finger position and moving the finger passively. In the Active condition, participants made back and forth movements between two bounds while their finger position was displayed as a green dot on the screen. During the course of the second movement, finger position became invisible and shortly after, a probe (blue dot) appeared while a brief tone was played. Participants responded if they thought the probe had appeared ahead or behind of their finger position and reported their confidence in their response. In the Passive condition, the task was identical except that the finger was moved by the robotic arm which reproduced a previous active movement. Participants could then use proprioceptive feedback and visuo-temporal cues to make up their mind. In the Visual condition, the task remained the same but the participants did not move their finger, a previous movement being only replayed on the screen. Therefore, the decision was then based solely on visuo-temporal cues of the movement.

## 2 Material & Methods

### 2.1 Participants

Twenty-nine right-handed participants were recruited (mean age = 22.62, SD = 2.7). The robotic device had limited power, so we selected participants with small hands. As a result, the majority (27/29) were female. Technical difficulties with the robotic arm prevented full testing of two participants. Their data were not analyzed. Two other participants were excluded as they presented strong response bias (responding *ahead* or *behind* in more than 75% of trials) that precluded meaningful signal detection analysis. Therefore, the final sample included 25 participants (24 female, mean age = 22.4, SD = 2.5). All participants had normal hearing, normal or corrected-to-normal vision, and no psychiatric or neurological history. They were naive to the purpose of the study and gave informed consent. The study was approved by the university ethical committee.

### 2.2 Movement task

Participants sat in front of a 525 × 320 mm computer monitor while their index finger was attached to a robotic device (Phantom Premium haptic device, Geomagic) able to record the finger position and actively move the finger.

Participants rested their right arm onto a support positioned parallel to their body at a comfortable height. The hand posture allowed the right index finger to make flexion and extension movements (Figure 1). The distal segment of their index finger was attached to the robot using a Velcro loop. Participants viewed the screen in front of them, and were instructed not to look directly at their hand. They were shown the position of their finger in the form a green dot of 4.3 mm diameter that they could move. On the screen, a white rectangular frame of 40 × 70 mm was presented, with top and bottom edges being bounded by a 7mm grey zone, and with a central red cross.

Participants were instructed to move between the two bounds of the frame at a constant speed, between 3.71 and 9.63 cm per second. Feedback on the velocity of the finger movement was given by the changes in colour of the green dot showing their finger position (blue = too slow, red = too fast). Participants had approximately three minutes to familiarize moving using this arrangement. After the training, participants were instructed to reproduce the same types of movement during the main experiment.

### 2.3 Trial procedure

After the training phase, participants were instructed regarding the main task, starting with the Active condition. At the beginning of each trial, participants moved the index finger to bring the green cursor onto the central red cross. An arrow indicated whether the first movement should be a flexion or extension. They then made movements back and forth between the bounds of the white frame. Each time their finger reached the bound of the frame, the bound changed from grey to yellow, indicating a change of direction was required.

Each trial involved three successive and continuous movements back and forth. During the first movement, from the center of the screen to the bound designated by the arrow, the green cursor continuously displayed the finger position. During the second movement, the green dot suddenly disappeared at a random location. The bound still changed from grey to yellow when touched, indicating when to change movement direction. During the third movement, a probe, represented by a blue dot of a diameter equal to the green dot appeared while a brief tone was played through headphones. Importantly, the probe appeared ahead or behind of the moving finger. Participants finished their movement, indicated by the last bound turning red. Then, the blue dot and frame disappeared and the words “Ahead” and “Behind” were displayed on each side of the screen. Participants responded to indicate whether the probe had appeared ahead or behind of their instantaneous finger position, by pressing one of two keys with the left hand. The response was unspeeded. Finally, the question “How confident are you in your response?” was displayed on the screen with the number 1 to 4 displayed underneath, 4 corresponding to maximal confidence. Initially, one random number was circled and participants moved the circle by pressing keys with the left hand, using a third key to register their confidence judgment.

To ensure that participants did not change the velocity of their movements, trials were interrupted when participants exceeded a speed of 16.96 cm per second. Trials could also be interrupted if people did not respect the imposed first movement direction or if they stopped moving too soon after the probe appeared. Participants were explicitly told that those interruptions were no errors but only means to improve their performance in the discrimination task.

The gap between the instantaneous finger position and the probe was adjusted to control task difficulty (see staircasing procedure). “Behind” and “Ahead” trials were randomly intermixed. Importantly, for both types of trials, the probe appeared at a random location chosen uniformly within the same central region of the frame, so that its position could not be used to predict the required response. This central region was defined so that the probe could never appear less distant to the bounds than the maximal gap distance recorded for that block.

### 2.4 Movement replay

Participants started the experiment with a training block of active trials, the 2-D coordinates of the position of the finger being recorded every millisecond. Next, they received instructions for passive trials. The passive trials followed the same procedure as the active trials, except that participants were instructed to keep their finger relaxed and avoid any voluntary movement. Instead, the robotic device reproduced a previous movement made by the participant. In order to check that no voluntary movement interfered with the robot’s command, movement’s trajectories with a velocity inferior or superior to 10% of the required velocity were stopped and the trial was restarted. As before, participants judged whether the probe was presented ahead or behind of their finger position, and reported their confidence in that judgment.

In the visual condition, participants were instructed to not move their finger at all. The trajectory of a previous active movement was replayed on the screen in a similar way, but the finger and device remained still. Participants had now to judge whether the probe was presented ahead or behind of the calculated position of the green dot, based only on visual-temporal cues such as the initial movement path displayed, and the colour change of the bounding zones.

### 2.5 Adaptive difficulty and experimental procedure

Because first-order performance have a strong impact on second-order metacognitive performance (Galvin et al., 2003; Maniscalco & Lau, 2012), we used a staircase procedure to equate performance between the active, passive and visual condition. To adjust the difficulty of the task in each condition, we varied on a trial-by-trial basis the gap between the probe and the actual finger position, larger distances making the task easier while smaller distances made the task more difficult. A 1-up/2-down staircase procedure was used to find the gap value eliciting 71% correct ahead/behind judgements (García-Pérez, 1998). Since moving objects are generally perceived ahead of their actual position (Nijhawan, 2001), the gap between the probe and the finger position was varied independently for “Ahead” and “Behind” trials, in two separate staircases. “Ahead” gaps were larger than “Behind” gaps for most participants, but accuracy for the two trial types was not affected by a bias towards one response.

We provided feedback for an initial 4-12 familiarization trials only, by showing both the actual finger position and the position of the probe (blue dot) at the end of each trial. Then, participants continued with two blocks of 40 trials to allow the ahead and behind staircases to converge to stable values. This procedure was repeated for each condition, starting with the active condition, then passive and then visual, taking approximately twenty minutes for each. During the main experiment, the staircase procedure was maintained, reducing the size of the incrementing steps (from eight pixels increments to five pixels increments during main experiment).

An experiment consisted in two sessions of 1 hour and a half each. Sessions were executed within the same day or on two consecutive days. The main experiment consisted in a total of fifteen blocks of thirty-six trials each, five blocks per condition. The order of the blocks was randomized within each session. Each active block was replayed in the passive and visual conditions in a random order. Within each block, active movements were replayed in a random order.

### l.l. Kinematic analyses

The velocity profile of the crucial third movement was retrieved from the robotic interface, and aligned to movement onset or to appearance of the probe. Velocity traces were averaged within each condition for each participant, then grand-averaged across participants for display. To determine statistical differences in velocity between the active and the passive conditions, a cluster-based non-parametric test with Monte Carlo randomization (adapted from Maris and Oostenveld, 2007) was applied. This method allowed us to identify clusters of time-points in which velocities differ, with appropriate correction for multiple comparisons.

In order to identify whether element of the movement influenced accuracy and/or confidence, we computed for each trial the mean velocity in the movement direction (y-axis) by averaging the velocity from the onset of the movement (first point after change of direction) to the reaching of the opposite bound. We also computed the lateral displacement of the movement, computing the distance from the most leftward point to the most rightward point of the trajectory.

### 2.6 Behaviour analysis

The first three blocks (training phase for each condition) were excluded from the analysis. Paired t-tests (two-tailed) and repeated measures ANOVAs were used to compare mean accuracy, mean gap values, mean RT and mean confidence. In order to quantify the support both in favor of accepting or rejecting the null hypothesis, we also computed Bayes Factor measure for the planned comparisons (Rouder, Speckman, Sun, Morey, & Iverson, 2009). We report BF01 which provides a measure of support towards the null hypothesis. In particular, values of BF01 > 3 provides positive support in favour of the null hypothesis while value BF01 < 0.1 provide positive support to reject the null hypothesis (Rouder et al., 2009).

To analyze metacognitive abilities, meta-d’ was computed with the Matlab toolbox provided by Maniscalco & Lau (Maniscalco & Lau, 2016). The meta-d’/d’ ratio was taken as a bias-free measure of metacognitive efficiency (Maniscalco & Lau, 2016). To analyse metacognitive bias across conditions, we retrieved the second-order criteria fitted for the computation of the meta-d’ for each participant and each condition. Note that as confidence was reported on a 4-point scale in the experiment, we obtained three separate criteria, for the 1-2, 2-3 and 3-4 boundaries respectively, independently for “ahead” and “behind” responses. We then computed absolute distance of the criteria to the first-order decision threshold for each response side, to provide a measure of how participants rated their confidence according to the level of internal evidence. For example, if a participant set the 3-4 confidence boundary very close to the first order decision criterion, then a very small degree of evidence would make them highly confident. As sensitivity and bias might vary across participants and conditions, we designed a normalization procedure which would allow us to determine how optimally participants’ confidence ratings tracked accuracy of their first-order decisions.

To do this, we developed a method to measure the optimal confidence criterion of each participant and estimate how they positioned their actual confidence criterion relative to that optimal criterion (Figure 2). For the meta-d’ value and decision threshold found for each participant, we simulated all the possible values of second-order hit (HIT2) and false-alarm (FA2) rates obtained by shifting a single criterion distinguishing low from high confidence trials (261 second-order criteria simulated) on the decision axis (Illustration of the process for one participant, Figure 2A-B). For each point, HIT2 and FA2 were calculated separately for each response i.e. each side of the decision threshold (c1), allowing to retrieve, according to second-order signal detection theory, the two full second-order receiving-observer curves (ROC2) associated with each response (Maniscalco & Lau, 2012). We then computed the subtraction HIT2 – FA2 (Figure 2, C-D) for each of simulated values and found the maximum of this difference, establishing the optimal second-order criterion (Figure 2, red circle) allowing to report high confidence with the highest hit rate and the lowest false-alarm rate (see Figure S1 for simulations of the optimal confidence criterion for different values of d’ and first-order criterion). We then retrieved how the criteria corresponding to each confidence rating boundaries (Figure 2, blue triangles) were positioned compared to this optimal criterion. To do so, we normalized the criterion values by the distance between the optimal second-order criterion and the first-order criterion so that this distance would correspond to a unit of one. Therefore, according to that measure, the zero value would correspond to the position of the first-order decision threshold (c1) and a value of one would correspond to the position of the optimal second-order decision threshold. This expresses how confidence criteria are placed on the decision axis in a way that is meaningful irrespective of first-order sensitivity and bias, though the method is potentially affected by the quality of the fitting of the meta-d’ quantity. For clarity, we averaged together criterion of each response side (“ahead”/“behind”), and transformed to values a logarithmic scale for statistical comparison.

**Figure 2:**
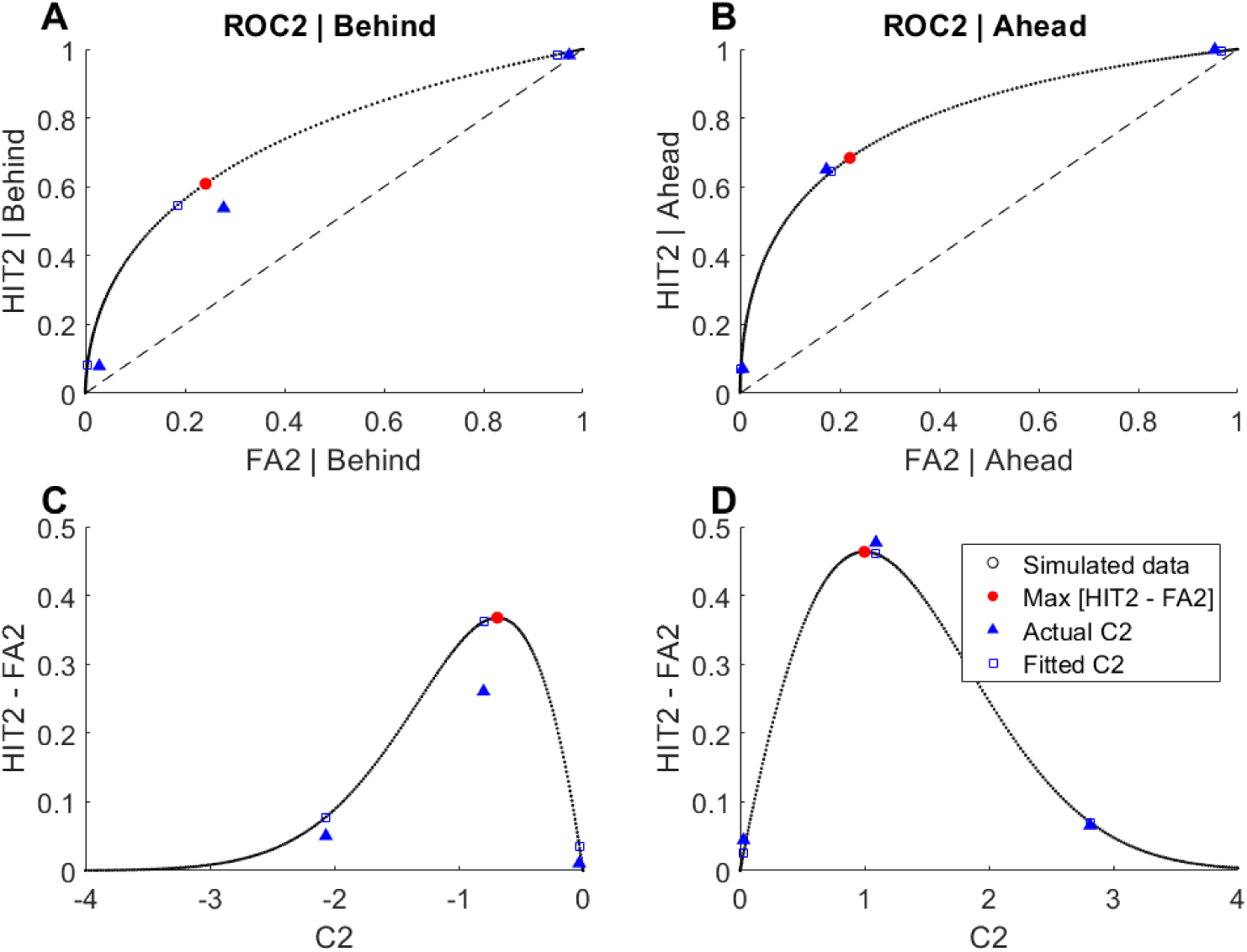
Illustration for one participant and one condition of the method used to retrieve optimal confidence criterion. For each participant and each condition, we computed for each response side (A,C: Behind; B,D: Ahead) the full second-order ROC2 curve corresponding to that first-order criterion and meta-d’ value (black dots). To do so, we varied along the decision-axis the position of a second-order “confidence criterion “ distinguishing low and high confidence trials and calculated the resulting proportions of second-order hits (HIT2 = p(High Confidence\Correct) and second-order false alarms (FA2 = p(High Confidence\Error). We then retrieved the difference between these HIT2 and FA2 rates (C-D) to find the second-order confidence criterion that maximized that difference. This value corresponds to the position of the second-order criterion (red dot) that allows to separate optimally error and correct trials for that particular value of meta-d’ and first-order bias. We used that “optimal confidence criterion” to normalize the values of actual criteria found for that participants (blue squares).

### 2.7 Predictors of accuracy and confidence

We investigated whether accuracy and confidence were influenced by the same factors and whether differences between the influence of these factors were observed across condition. To do so, we used multiple linear regression performed separately for each participant and each condition to determine the parameters that influenced response choice (ahead/behind), accuracy, and confidence. The regressors used, and a justification of their inclusion, are given in supplementary table 1. As some of these predictors were collinear (for instance the finger and probe position were r), we used a least absolute shrinkage and selection operator (LASSO, Tibshirani, 1996) regression which selects predictors and regularizes the linear model by assigning null values to redundant predictors.

To determine which factors influenced response choice, accuracy and confidence while avoiding instability in the estimation of the linear model, we transformed categorical predictors into continuous variables and used a bootstrapping approach. To do so, we randomly partitioned the total number of trials into 18 subsets of 10 trials and computed the average value of each regressor for each subset, allowing to obtain continuous values for each parameter (Wokke, Cleeremans, & Ridderinkhof, 2017). We then estimated the best linear model using LASSO regression and retrieved the beta values associated with each predictor for that model. We repeated this procedure 1000 times based on different random partitions of the data. We then computed the mean beta value over all repetitions and divided by the standard deviation of the beta distribution to obtain one beta estimate for each parameter and each participant (normalized beta value). We then tested whether the betas associated with each predictor differed from 0 across participants using a t-test approach.

## 3 Results

### 3.1 Accuracy, task difficulty & Confidence

The goal of the present experiment was to explore the contribution of voluntary motor command and proprioceptive information in motor awareness and metacognitive judgments. To do so we used a planned comparison approach, contrasting judgments on active and passive movements to determine the contribution of motor command to movement perception and comparing judgments on passive movements and visual trajectories to test the contribution of proprioceptive information to movement perception.

We first investigated whether our manipulation to equate performance across conditions was successful. This was achieved by using a 2down-1up staircase procedure adjusting the gap distance between the probe and the actual finger position (see Methods), smaller distances increasing the difficulty of the task. Although no large differences in accuracy were observed between conditions, accuracy remained significantly higher in the Active condition than in the Passive (Figure 3A, t(24) = 4.1 p < 10^-4^, d = 0.8) and in the Visual condition (t(24) = 2.67 p = 0.014, d = 0.53). This was observed despite the fact that the gap distance between the probe and the actual finger position reached by the staircase was significantly smaller in the Active condition compared to the Visual condition (Figure 3B, t(24) = -3.2 p < 10^-3^, d = -0.64) and to the Passive condition (t(24) = -1.95 p = 0.06, d = -0.39). The gap did not significantly differ between the Passive and Visual condition (t(24) = -0.945 p = 0.35, d = -0.19).

**Figure 3:**
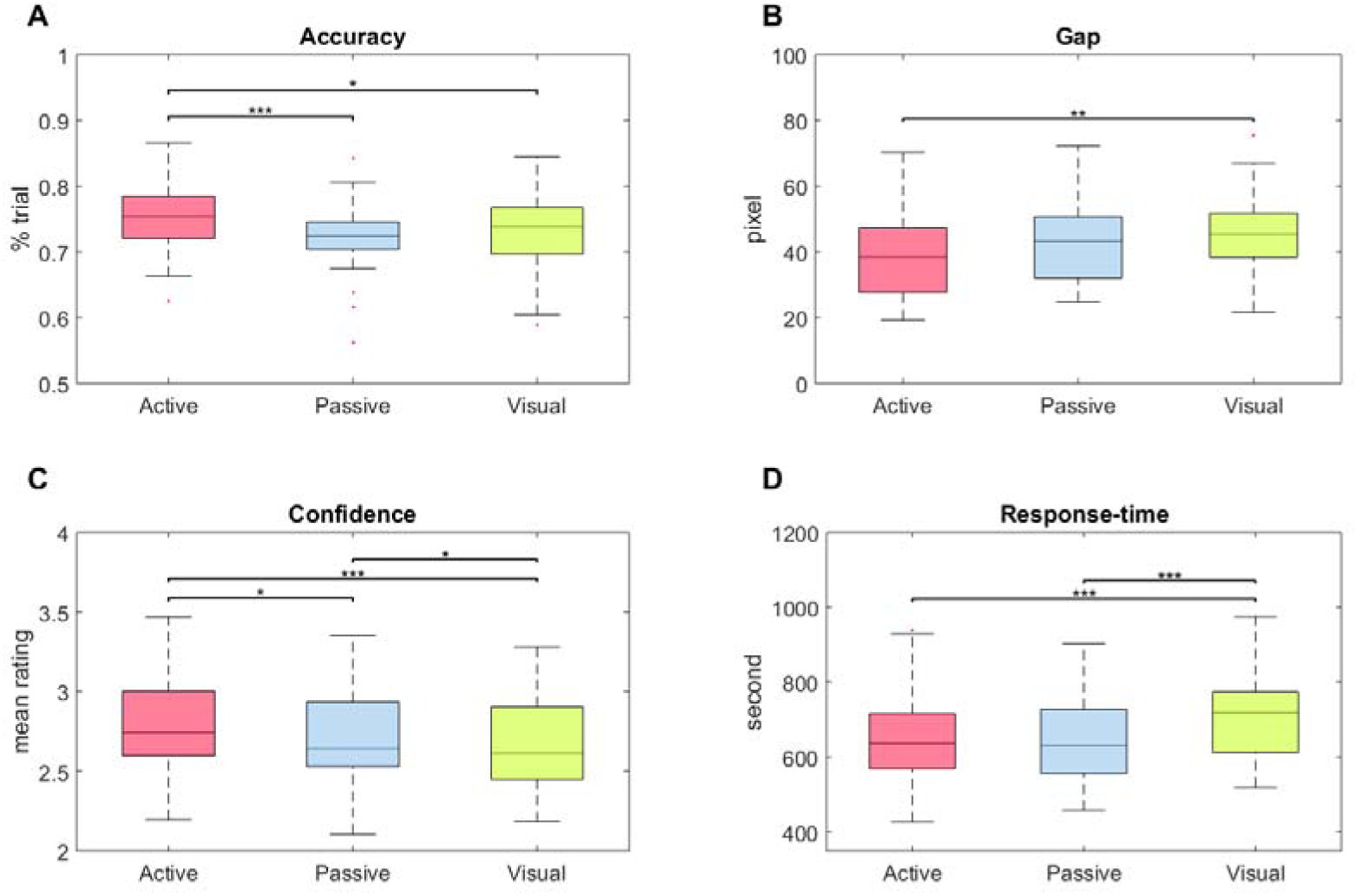
Boxplot of Accuracy, Gap (probe-finger distance), Confidence and Response time. A: Percentage of correct responses in the Active (red), Passive (blue) and Visual (green) conditions across trials and participants. B: Gap distance between the position of the probe and the actual finger position. Gap value was adjusted on a trial-by trial basis following a staircase procedure to equate decision accuracy between conditions. Smaller gap values indicate increased task difficulty. C: Confidence ratings (1-4 scale) for each conditions, across trials and participants. D: Response-time for each conditions, across trials and participants. For all plots, central mark indicates the median, and the bottom and top edges of the box indicate the 25th and 75th percentiles. Top black bars indicate significant difference with p <0.05:^∗^, p <0.01:^∗∗^, p <0.001:^∗∗∗^.

Average confidence followed the pattern of accuracy, participants being significantly more confident in the Active compared to the Passive condition (Figure 3C, t(24) = 2.61 p = 0.015, d = 0.52) and Visual condition (t(24) = 3.98 p < 10e-4, d = 0.8). Interestingly, participants also expressed higher confidence in the Passive than in the Visual condition (t(24) = 2.37 p = 0.026, d = 0.47) although no difference in accuracy was observed between these two conditions.

Response-time (RT) were overall slower in the Visual than in the Active (Figure 3D, t(24) = - 3.97 p < 10e-4, d = -0.79) and in the Passive condition (t(24) = -3.79 p < 10e-4, d = -0.76) while no significant difference in RT was observed between the Active and Passive condition (t(24) = -0.101 p = 0.92, d = -0.02).

Taken together these results confirmed participants’ performance increased from the Visual condition to the Active condition. Additional analysis (see Supplementary Results 2.1 and Figure S2-3) revealed that these differences they were not due to voluntary change in the movement in the Active condition but to a better estimation of the finger trajectory when based on voluntarily motor command than on proprioceptive feedback or visuo-temporal cues alone.

### 3.2 Second-order signal detection analysis

#### 3.2.1 First and second-order sensitivity

To evaluate potential metacognitive differences between conditions, second-order signal-detection theory method was used to compute d’ and meta-d’ values for first-order and second-order sensitivity (see Methods). Response and confidence bias were also estimated. D’ and meta-d’ measures are independent of each other and the ratio between them provides an estimate of metacognitive efficiency, controlling for effect of first-order accuracy and confidence bias (see Methods).

Measure of d’ (first-order sensitivity, Figure 4A) followed the same pattern as accuracy, suggesting that participants remained significantly better in the Active condition than in the Passive condition (t(24) = 3.54, p < 10e-3, d = 0.71, BF01 = 0.05) and the Visual condition (t(24) = 2.54, p = 0.018, d = 0.51, BF01 = 0.41) despite the staircase procedure. No significant difference in d’ was observed between the Passive and the Visual conditions (t(24) = -1.02, p = 0.32, d = -0.2, BF01 = 4).

**Figure 4:**
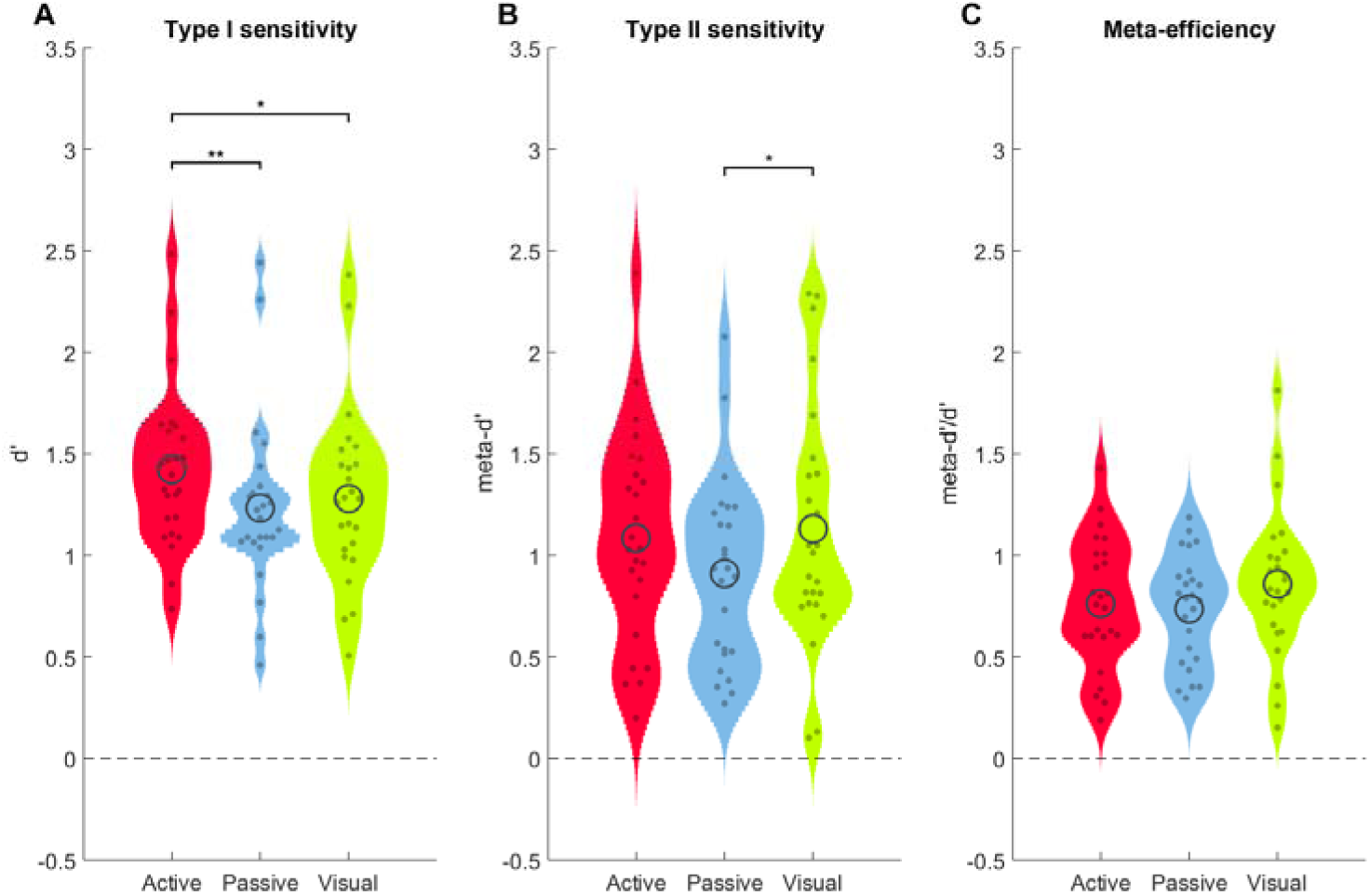
First-order sensitivity, second-order sensitivity and metacognitive efficiency. Violin plot of d’ measures (A,) meta-d’ measures (B) and meta-d’/d’ ratio (C) across participants for Active (red), Passive (blue) and Visual (green) conditions. Full dots represent individual values. Black circle represents the population mean. Top black bars indicate significant difference with p <0.05:^∗^, p <0.01:^∗∗^, p <0.001:^∗∗∗^.

At the second-order level, meta-d’ (second-order sensitivity, Figure 4B) revealed no significant difference between the Active and the Passive conditions (t(24) = 1.67, p = 0.11, d = 0.33, BF01 = 1.8) or between the Active and Visual condition (t(24) = -0.448, p = 0.66, d = -0.09, BF01 = 5.9). However, a significant difference was observed between the Passive and Visual conditions (t(24) = -2.24, p = 0.034, d = -0.45, BF01 = 0.71).

As such result could be the result of the observed differences in first-order performance, we turned to the ratio of meta-d’/d’ (Figure 4C). There were no significant differences however between conditions in this measure of metacognitive efficiency, neither between Active and Passive conditions (t(24) = 0.405, p = 0.69, d = 0.081, BF01 = 6), nor between Active and Visual conditions (t(24) = -1.45, p = 0.16, d = -0.29, BF01 = 2.4) or between Passive and Visual conditions (t(24) = -1.48, p = 0.15, d = -0.3, BF01 = 2.4).

Overall, these results show that when making a judgment on the position of a moving object, whether simply observing the movement, being moved passively or making the movement voluntarily, no difference was observed in metacognitive abilities once task difficulty was properly controlled.

#### 3.2.2 Correlation between modalitis in first and second-order sensitivity

We further investigated whether first- and second-order sensitivity correlated between conditions, potentially suggesting a common factor underlying perceptual and metacognitive judgements in all three conditions (see Figure S4 and table S1 for full results). We found that d’ correlated significantly between all conditions (all p < 10-3), as did meta-d’ (all p < 0.02). Metacognitive efficiency correlated between the Passive and Active conditions (p = 0.028) as well as between the Visual and Active conditions (p < 0.01) but not between the Visual and Passive conditions (p=0.45). Taken together, these results suggest that both first- and second-order performance were related between tasks although the correlation did not reach significance between the Passive and the Visual conditions.

#### 3.2.3 Difference in confidence bias between conditions

Next we explored how participants set their decision and confidence criterion in each condition.

At the first-order level, no bias towards “Ahead” or “Behind” responses were observed, first-order decision criterion being centred on 0 in all the conditions (see Figure S5 and corresponding paragraph in the supplementary results). Furthermore, we found a significant correlation in the first-order decision threshold between each pair of conditions (all p < 0.001) suggesting that biases in decision threshold were shared between Active, Passive and Visual tasks (Figure S9).

Turning to potential biases in confidence ratings, we first estimated raw confidence ratings in error and correct trials in each condition (Figure S6). We found that average confidence in error and correct trials differed across conditions: participants were more confident in their correct responses in the Active than in the Passive (t(24) = 2.61, p = 0.015, d = 0.52, BF01 = 0.35) and in the Visual (t(24) = 3.65, p < 10e-3, d = 0.73, BF01 = 0.038) conditions. Conversely, they were less confident when they actually made an error in the Visual compared to the Active (t(24) = 3.27, p < 10e-3, d = 0.65, BF01 = 0.09) and the Passive (t(24) = 3.35, p < 10e-3, d = 0.67, BF01 = 0.075) conditions. As no differences in metacognitive efficiency were observed between those conditions, we expected these differences to result from a change in confidence bias across conditions.

Second-order signal detection theory proposes that different levels of confidence is obtained by placing additional second-order criteria on either side of the first-order decision criterion. If the evidence falls close to the first-order decision boundary, the confidence in the response will be rated as low. If on the other hand the evidence falls farther from the decision boundary, the response will be labelled as made with high confidence (Maniscalco & Lau, 2012, 2016). A similar model can be used when confidence is not just rated as High or Low but with graded levels, as in the present study. In that case, one criterion is fitted for each boundary between confidence ratings (see also figure 5).

**Figure 5:**
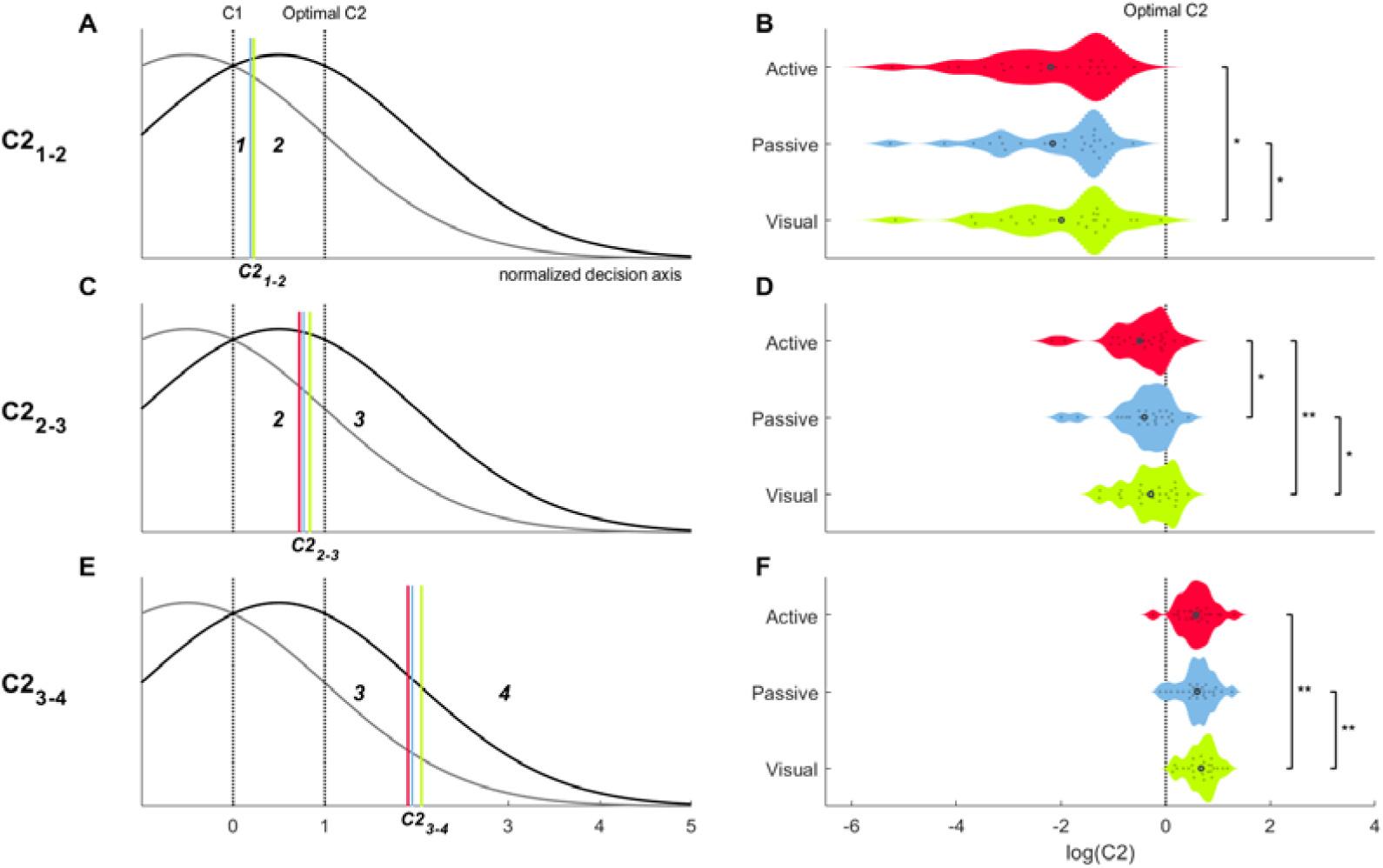
Type II criteria. Mean position of the normalized second-order criteria on the decision axis (A,C,E) and violin plot their distribution on a logarithmic scale (B,D,F) across participants for Active (red), Passive (blue) and Visual (green) conditions. Type II criteria were retrieved from meta-d’ fitting procedure for the boundary between each confidence ratings. Their distance to the decision criterion (C1) was then calculated for each response side separately. This distance was normalized by dividing it by the optimal decision criterion The normalized distance of these criteria were averaged together across response side using absolute value. The first column (A,C,E) represents a schematic of the mean position of each criterion the decision axis for each condition. The second column (A,C,E) shows the violin plot of the corresponding distributions, values being transformed using logarithmic scale. Full dots represent individual values. Black circle represents the population mean. Vertical black bars indicate significant difference with p <0.05:^∗^, p <0.01:^∗∗^, p <0.001:^∗∗∗^.

To analyse differences in how confidence criteria were set among conditions, we retrieved the second-order criteria fitted for the computation of the meta-d’ for each confidence rating boundary and calculated their absolute distance to the decision threshold for each response side. In order to understand how the criteria were positioned on the decision axis, we compared those values to the position of an optimal confidence criterion calculated for each participant and each condition according to their meta-d’ and first-order decision criteria. This optimal criterion was defined as the criterion value allowing for the greater difference between the proportion of Correct trials associated with high confidence (HIT2) and lower proportion of Errors associated with high confidence (FA2) (See Methods). We use that value to normalize participant’s second-order criteria, allowing us to compute a measure of criterion shift independent of both first-order accuracy and first-order criterion. For clarity, we averaged both response side (“ahead”/“behind”) together and used a logarithmic scale to assess differences between conditions (an analysis of the criteria before normalization can be found in supplementary material, Figure S7).

We found that the boundary between confidence ratings 2 and 3 was placed the closest to the optimal confidence criterion (corresponding to a value of 1 on Figure 5A,C,E and a value of 0 on the logarithmic scale on Figure 5B,D,F), suggesting that participants placed the separation between Error and Correct trials close to the middle of the confidence scale. Nonetheless, the intermediate criterion did significantly differ from the optimal criterion in all conditions (Figure 5; Active: t(24) = -4.16, p < 10e-4, d = -0.83, BF01 = 0.012; Passive: t(24) = -3.76, p < 10e-4, d = -0.75, BF01 = 0.03; Visual t(24) = -3.08, p < 10e-3, d = -0.62, BF01 = 0.14) suggesting that participants placed the boundary between perceived Error and Correct response toward lower confidence ratings rather than the exact middle of the scale. That is, participants required surprisingly less than expected evidence to report above-median levels of confidence.

Moreover, we also observed that the position of the criteria was different across conditions. Overall, criteria were positioned closest to the decision threshold for the Active condition, followed by the Passive and then the Visual condition. A significance difference was observed between the Visual compared to the Active and Passive conditions in the position of the lowest confidence criterion (boundaries between confidence 1 and 2: Active vs Visual t(24) = -2.24, p = 0.017, d = -0.45, BF01 = 0.72; Passive vs Visual t(24) = -2.1, p = 0.023, d = -0.42, BF01 = 0.91) and the highest confidence criterion (Active vs t(24) = -2.88, p < 10e-3, d = -0.58, BF01 = 0.2; Passive vs Visual t(24) = -2.98, p < 10e-3, d = -0.60, BF01 = 0.17). For the intermediate criterion corresponding to the limit between confidence ratings of 2 and 3, a significant difference was observed between the three conditions (Active vs Passive t(24) = -2.09, p = 0.024, d = -0.42, BF01 = 0.92;Active vs Visual t(24) = -3.38, p < 10e-3, d = -0.68, BF01 = 0.071;Passive vs Visual t(24) = -2.17, p = 0.02, d = -0.43, BF01 = 0.81).

Taken together, these results suggest that participants were progressively more liberal in their confidence judgments across conditions: at equal levels of evidence for a first-order decision, they were significantly more likely to give higher confidence ratings in the Active than in the Passive condition, and in the Passive than the Visual condition. Additional analysis confirmed this shift in criterion was not entirely explained by a change in first-order accuracy (Figure S8) as no correlation was observed between the difference in second-order criteria and the change in accuracy between conditions. Interestingly however, a positive correlation was found in the average position of the second-order criteria across conditions, suggesting that some common process underlay confidence rating across conditions (Figure S9).

#### 3.2.4 Factors influencing accuracy and confidence

Finally, we wanted to shed some light on the factors that influenced first-level performance and second-level metacognition in each condition. To do so, we used multiple linear regression performed separately for each participant to determine the parameters that influenced accuracy and confidence. The list of regressors, and a rational for their inclusion, is shown in table 1. Because of possible redundancy and multicollinearity between regressors, we used a LASSO regression approach (Tibshirani, 1996) which sets to 0 redundant predictors, therefore reducing effect of collinearity.

**Table 1:** List of regressors included to predict response choice, decision accuracy and confidence

| VARIABLE NAME | TYPE | DESCRIPTION | JUSTIFICATION |
| --- | --- | --- | --- |
| *Response* | Categorical | Response made by the participant: "Behind" or "Ahead" | Participants might be more accurate/confident for one of the response options |
| *Behind or Ahead* | Categorical | Probe being ahead or behind of the finger. | Accuracy might differ for Behind and Ahead trials. |
| *Probe position* | Continuous | Position of the probe on the screen relative to the onset of movement. | Participants might be more accurate for some positions of the probe than others |
| *Finger position* | Continuous | Position of the finger when the probe appeared relative to the onset of movement | Participants might be more accurate to estimate their finger position at certain phases of the movement |
| *RT* | Continuous | Time taken to respond after the presentation of the response options on the screen | Response-time might correlate with Accuracy and Confidence |
| *Gap* | Continuous | Distance between the finger position and the probe | Participants should be more accurate for larger gap distance |
| *Probe distance to centre* | Continuous | Distance of the probe relative to the centre | Participants might be more confident when the probe appear closer to the bounds. |
| *Finger distance to centre* | Continuous | Distance of the finger to the centre at the time of the apparition of the probe | Participants might be more confident when their finger is close to the bounds. |
| *Flexion or Extension* | Categorical | Movement corresponded to a flexion or an extension of the finger | Accuracy might be affected by the direction of movements |
| *Velocity (y-axis)* | Continuous | Mean velocity in the direction of the movement (Velocity y-axis, (see Supplementary Methods) | Increased velocity might increase difficulty in position judgment. |
| *Displacement*<br>*x-axis* | Continuous | Displacement to the direction perpendicular<br>to the direction of the movement (see<br>Supplementary Methods) | Larger lateral movements might<br>lead to poor position estimation<br>along the principal movement<br>axis |

As an initial sanity check, we first considered which factors predicted “ahead” vs “behind” response choice (Figure 6A). As might be expected, the relative position of the probe compared to the finger correlated with response choice, explaining more variance than the actual correct response (Ahead or Behind). More surprisingly, longer RTs were associated with “behind” responses, suggesting that inattention or difficulty in responding were associated with poor predictive representation of hand position.

**Figure 6:**
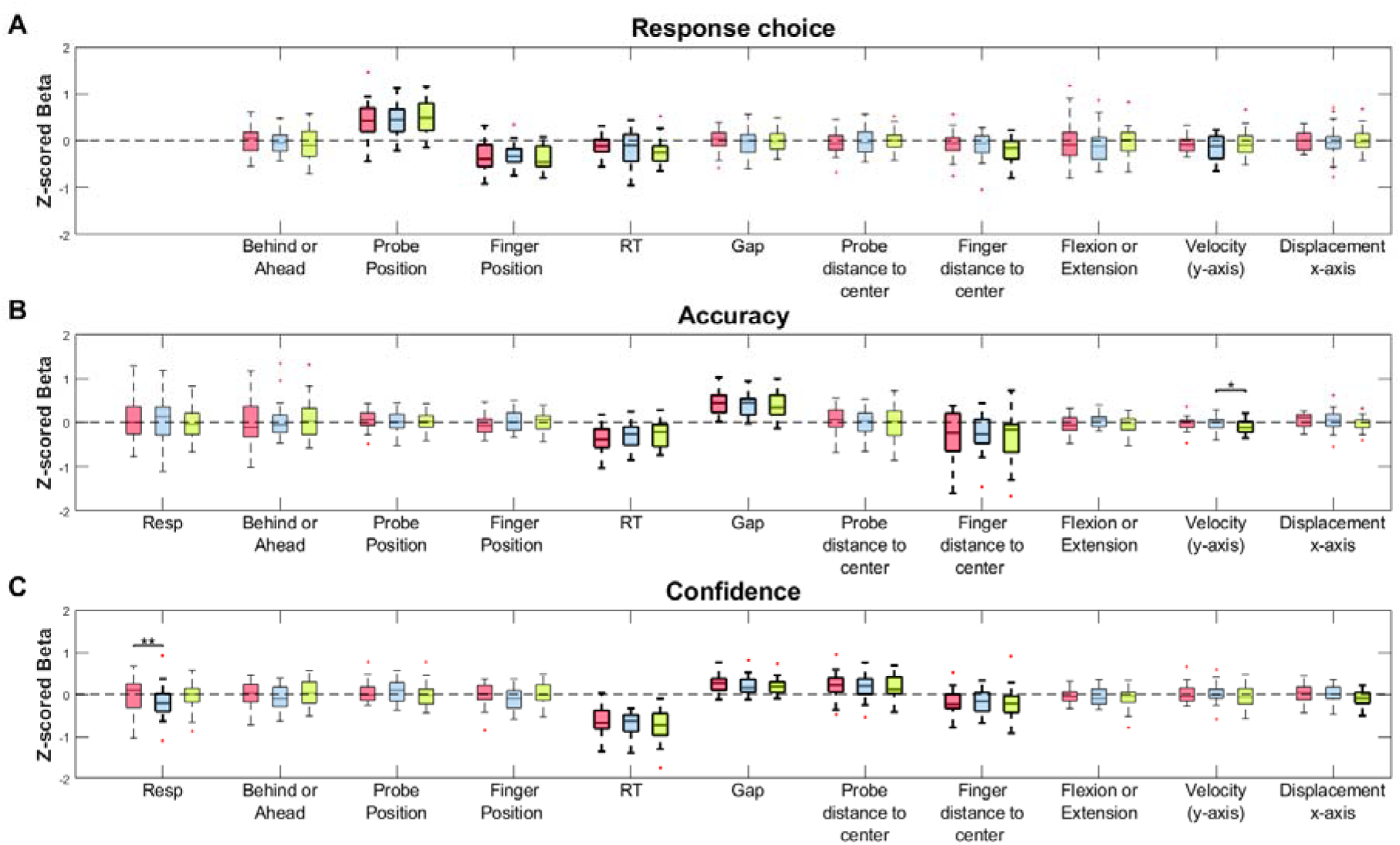
Boxplot of the beta coefficients of the multiple linear regression predicting Response Choice (A), Accuracy (B) and Confidence (C) for the Active (red), Passive (blue) and Visual (green) conditions. Each multiple regression was performed separately for each condition and each participant. A bootstrapping approach was used to estimate the beta coefficient of each predictor: random partitioning of the trials and calculating of the regression coefficients was repeated 1000 times. The obtained betas coefficients were then z-scored across repetitions and then average across participants. We then tested whether the obtained z-score beta coefficient differed from 0 across participants, significant boxplot being displayed in bold. Central marks represent the median value of the obtained coefficient across participants, while top and bottom edge represent the 25^th^ and 75^th^ percentile. Whiskers represent most extreme values and outliers are displayed as red crosses.

We next explored predictors of decision accuracy (Figure 6B). First, we found that RT correlated with accuracy, more errors being committed for longer RTs, as might be expected. Unsurprisingly, accuracy was also predicted by the distance between the probe and the finger position (Gap), larger gaps predicting more correct responses. More surprisingly, we found that the closer the finger was from the bound of the box (Finger distance to centre), the more participants made errors. This result is surprising as the required response was actually more predictable when the finger was closer to the bound, making the task easier for those trials.

Our main interest lay in how the same model explained confidence judgments (Figure 6C). We found that confidence decreased with longer RT. Interestingly however, beta values were significantly higher than for accuracy (t-test for each condition, all p < 10^-3^), suggesting a stronger impact of RT on confidence. We found that larger gap values correlated with higher confidence but the beta values were significantly lower than for accuracy (t-test for each condition, all p < 10^-3^). Regarding the impact of finger position, confidence followed the pattern of accuracy, being significantly lower when the finger was more distant to the centre. This suggests that participants were aware that they were making more mistakes for trials in which the finger was far from the centre, this factor having a similar impact on confidence and on accuracy (t-test for each condition, all p > 0.08). Surprisingly however, we found that participants reported stronger confidence when the *probe* appeared farther from the centre, although this predictor did not correlate with accuracy. This result seems to suggest that participants made false assumptions about the difficulty of the decision according to the position of the probe.

Overall, these analyses showed that many factors influencing response accuracy also influenced confidence, confirming participants were at least partially aware of what caused them to make errors. Interestingly, some parameters seemed to impact only confidence, reflecting incorrect beliefs influencing the difficulty of the task. In particular, a purely visual feature of our probe task which was unrelated to actual perceptual performance had a significant influence on confidence suggesting a form of metacognitive ‘hallucination’. We speculate that the visually salient event of a highly eccentric probe lead to a high confidence, even though this visual information was irrelevant to the task. Importantly, no significant differences were found across conditions on how these parameters influenced accuracy and confidence.

## 4 Discussion

In the present study, we investigated the metacognitive abilities related to voluntary actions and passive movement perception, and a baseline condition involving visual information only. Our systematic study revealed several novel findings. First, although the accuracy of first-order decisions increased slightly for voluntary compared to passive movements and visual perception, no differences in metacognitive efficiency was observed between tasks when controlling for these variations in first-order accuracy. Second, metacognitive sensitivity and bias in confidence judgments were correlated between tasks across individuals, suggesting that a common process underlay metacognitive judgment for voluntary actions, passive movement and for purely visual decisions. Third, our results revealed that participants were more biased towards higher confidence ratings when judging their own voluntary movements then when judging movements executed passively, or when judging a visual replay of their movement. This result suggests an element of over-confidence when making judgements about one’s own actions. Finally, regression analyses suggested that participants had partially wrong beliefs about the factors influencing their accuracy, and used irrelevant task parameters as proxies when giving confidence ratings. Taken together, these results suggest that confidence judgements about voluntary actions involve biased estimates of accuracy.

The main objective of the present study was to determine whether there were differences in metacognitive abilities when judging voluntary movements, passive displacement of the limbs or when making decision about the movement of visual objects. We did not find differences in metacognitive sensitivity associated with these three types of judgment. Accuracy and metacognitive efficiency correlated strongly across tasks, recalling recent findings of a correlation in metacognitive judgment across sensory modalities (Faivre et al., 2017; Song, Schwarzkopf, Kanai, & Rees, 2011) or between types of decisions (McCurdy et al., 2013). Such a result is compatible with the view that metacognition constitutes a supra-modal process, extending these findings to proprioceptive and voluntary movement judgments. Thus, confidence and error detection in action and perception rely on a common cognitive function (Fleming et al., 2010), suggesting that confidence signals act as a common currency measure to evaluate and compare performance across tasks (Ais et al., 2016; De Gardelle et al., 2016).

Could an alternative hypothesis explain the absence of differences in metacognitive sensitivity between the three tasks? One possibility is that the similarities at the metacognitive level are due to the similarities of the task in the three conditions. Indeed, all decisions required to judge the position of a probe compared the position of a moving object, relying either exclusively on temporal and visual cues, proprioceptive feedback or voluntary motor command. As all movements were replays of movements executed previously by the participant, it is therefore possible that participants relied on motor predictions in all three conditions to judge the relative position of the probe. Another alternative hypothesis is that metacognitive sensitivity differs between action perception and exteroception only when judging the overall success of the action, rather than the actual spatial path of the movement. Indeed, it has been proposed that motor awareness is dominated by representation of the goal of the action rather than representing the actual movement trajectory (Blakemore & Frith, 2003; Blakemore et al., 2002). Therefore, it is possible that, despite the results presented here, metacognitive sensitivity is increased when monitoring action success compared to spatial path.

While further studies will be necessary to assess the fine contribution of motor predictions in metacognition of action, our findings confirm its importance in motor awareness. Performance was significantly increased when judging voluntary actions, despite our efforts to equate accuracy between conditions. In that respect, our result seems in accordance with the findings of a previous study showing that movement perception is improved for active compared to passive movements (Farrer, Franck, Paillard, & Jeannerod, 2003; Paillard & Brouchon, 1968). These results seem to confirm the importance of the efferent copy in action perception (Blakemore & Frith, 2003; Blakemore et al., 2002) demonstrating that motor predictions improve the representation of movements.

Despite not observing a difference in metacognitive sensitivity, we observed a difference in confidence bias across conditions. Overall, we found that participants tended to be more confident when judging their own voluntary actions than when judging passive finger displacement or visual trajectories of their own movements, placing their confidence criterion closer to the decision threshold. Importantly, this result did not appear to be only a consequence of the pattern of performance across conditions as the effect was observed when normalizing shift in confidence by an estimate of the optimal positioning of the criterion for that condition and that participant and the change in confidence criterion did not correlate with the increase in performance.

These analyses depend on individuals’ use of the confidence scale provided, so should be interpreted with caution. Nevertheless, we found that participants tended to be overconfident when judging their voluntary actions. What could be the basis of this bias? One possibility is that participants judged a priori that the Active condition was easier than the others, shifting their overall confidence towards higher ratings. Indeed, as the architecture of the task corresponded to an additive design, more information being gradually available from the Visual condition to the Active condition, participants might have make the corresponding prediction that they were performing gradually better in each condition. However, our finding that the shift of criterion did not correlate with the increase in performance (Figure S8) suggests that this hypothesis does not fully account for our results. An alternative account of these findings could be that this shift in criterion reflects a specific bias in confidence when judging our own movement and voluntary action. In that sense, it could echo the known overconfidence bias in introspective abilities, people believing they are better judge of their own actions than external observers (Jones & Nisbett, 1972; Nisbett & Wilson, 1977). An illusion of a privileged access to the information guiding our behaviour and the preeminence of intentions in perceiving our actions is thought be one of the cause of illusory perception of control over external events (Wegner, 2004) as well as of the illusory increased self-agency caused by subliminal priming (Moore, Wegner, & Haggard, 2009). This phenomenon of “apparent mental causation” can be linked to the “intentional binding” phenomenon which makes participants experience the consequences of their voluntary actions as happening sooner in time than normal (Kühn, Brass, & Haggard, 2013). In that respect, our finding of a confidence bias for voluntary action compared to exteroception fits with the view that volition potentially distorts action perception.

Finally, the present study also shed some lights on the factors influencing decision accuracy and confidence. Unsurprisingly, we found that both accuracy and confidence were influenced by parameters related to task difficulty, in particular the gap distance between the probe and the finger position, and the time taken to make a response, showing that participants were at least partially aware of the difficulty of the decision to make and its consequence on their response choice. Furthermore, confidence also correctly reflected some other parameters influencing decision accuracy such as the position of the finger at the time of the apparition of the probe. Interestingly however, confidence also varied with some parameters that did not actually impacted accuracy: participants reported higher confidence when the probe appeared further from the center although they did not appear to be more correct for those trials. Such finding speaks in favour of a dissociation between choice and confidence, suggesting some visual cues altered confidence specifically. This result is of particular interest as it shows that some irrelevant information can impact confidence, in accordance with findings that confidence does not simply reflect the continued processing of the same evidence that influenced the choice (Resulaj, Kiani, Wolpert, & Shadlen, 2009; van den Berg et al., 2016) but might incorporate distinct information and beliefs about decision accuracy and task difficulty (Navajas et al., 2017).

Taken together, the results of the present study shed new light on the awareness of actions. Our result provides the first investigation of the metacognitive process related to judging our own movement. It demonstrates that despite feeling more confident when judging our own voluntary movement, metacognitive processing of one’s own action is no more sensitive to first-order processing than metacognitive processing of exteroceptive signals. Our findings contribute to the understanding of metacognition more generally, and open new avenues of research in understanding how people perceive their own actions.

## Acknowledgement

This work was supported by a European Research Council Advanced Grant (HUMVOL, agreement number 323943), a Chaire Blaise Pascal of the Région ÎIle-de-France to PH and a post-doctoral fellowship of the British Academy to LH. The authors would like to thank Pascal Mamassian for his helpful comments on the design of the experiment.

